# Assessing local adaptation and divergence at early life stages within Eastern Baltic cod

**DOI:** 10.64898/2026.01.20.700346

**Authors:** Maddi Garate-Olaizola, Johanna Fröjd, Vanda Larsson Åberg, Alma Hodzic-Vázquez, Yvette Heimbrand, Anders Nissling, Jane W. Behrens, Maria Cortázar-Chinarro, Ulf Bergström, Anssi Laurila

## Abstract

Many commercially exploited fish stocks have declined over the last few decades. It is therefore essential to identify natural populations and understand local adaptation for sustainable management. Salinity is a key environmental factor shaping local adaptation, and adaptive trait divergence often occurs at the egg and larval stages. The strong salinity gradient in the brackish Baltic Sea has driven rapid adaptation in multiple taxa. The Eastern Baltic cod (*Gadus morhua*) has adapted to low salinity with buoyant and tolerant eggs and larvae, but the stock has declined both in abundance and geographical range during the last decades. The main reproduction area of this stock is in the Bornholm Basin (ICES subdivision (SD) 25) in the southern Baltic Proper. Cod in this area, however, exhibit stunted growth and small body sizes. In contrast, large and healthy cod in reproductive condition have been observed in the Åland Sea in the northern Baltic Proper (SD 29), raising the question of whether these fish represent a locally adapted population capable of successful reproduction in the lower salinities (5-10 psu in the northern Baltic Proper (SD 27, 29 and 32). Here, we experimentally assessed egg and yolk-sac larvae survival across salinities, egg size, egg and larval neutral buoyancy and egg survival on sediment, to test whether northern (Åland) cod show adaptation to low salinity at early life stages as compared to southern cod. Mortality of larvae increased with decreasing salinity in cod from both areas, with the lowest survival at 7 psu. At 9 psu, more than 50% of northern cod larvae survived, suggesting that development could occur in SD29. Egg size and buoyancy were similar between northern and southern cod, and eggs and larvae were negatively buoyant, sinking under local salinity conditions. Nevertheless, the eggs survived and hatched well on sediment, indicating potential for demersal spawning. Our findings show no strong evidence of adaptive divergence to lower salinity in northern cod; however, their ability to tolerate sediment contact at early life stages suggests that Eastern Baltic cod may reproduce outside their historical spawning grounds.

## Introduction

Many commercially exploited fishes have shown rapid population declines, caused by high fishing pressure in combination with habitat loss in recent decades (Worm et al. 2009; Brophy et al. 2020). This urges effective management to reduce exploitation and allow threatened populations to recover (Worm et al. 2009; van Gemert and Andersen 2018; Brophy et al. 2020). Identifying the appropriate management units or stocks for these populations is fundamental for effective management of fish; however, stock units often do not match natural populations (Hauser and Carvalho 2008; Hutchinson 2008; Brophy et al. 2020) and ignore local adaptation (Conover et al. 2006; Grabowski et al. 2011; Bonanomi et al. 2015; Momigliano et al. 2019), leading to mismatches in managing fishing pressure at the stock level. Adaptive divergence can occur at egg and larval stages and be essential for early survival and recruitment (Nissling et al. 1994; Momigliano et al. 2017; Nissling et al. 2017). Understanding adaptive divergence at these early life stages can therefore be crucial for effective stock management (Jokinen et al. 2019).

Environmental salinity is a crucial parameter for eggs and larvae of marine fish (Holliday 1969). Salinity influences egg buoyancy and determines where in the water column eggs will develop (Nissling and Westin 1991a; Nissling et al. 1994; Sundby and Kristiansen 2015; Nissling et al. 2017). More saline water has a higher density, and depending on its relationship with the specific density of the eggs, eggs can be positively buoyant and float or negatively buoyant and sink (Govoni and Forward Jr 2008). When the eggs match the salinity of the surrounding water, they reach neutral buoyancy and can remain in the water column (Govoni and Forward Jr 2008). These pelagic eggs are expected to die when they come in contact with the sediment on the sea floor (Hinrichsen et al. 2016). On the other hand, negatively buoyant eggs of demersal spawners sink to the bottom and tolerate being in contact with sediment (Lønning et al. 1988; Suthers and Frank 1991). Newly hatched larvae typically exhibit buoyancy characteristics similar to those of the eggs from which they originated (Davenport et al. 1981; Saborido-Rey et al. 2003; Govoni and Forward Jr 2008).

The Baltic Sea is a young (8000 yrs) semi-enclosed brackish sea (Leppäranta and Myrberg 2009). Across the Baltic Proper (ICES subdivision (SD) 25–29), salinity decreases sharply from approximately 18 to 5 psu along a south–north gradient, creating a strong selective environment for marine organisms (Pereyra et al. 2009; Johannesson et al. 2011; Wennerström et al. 2013). This has resulted in the emergence of unique lineages of several marine fish species adapted to the brackish Baltic environment, showing strong genetic divergence from the Atlantic populations (Limborg et al. 2009; André et al.. 2011; Lamichhaney et al. 2012; DeFaveri and Merilä 2013; DeFaveri et al. 2013; Berg et al. 2015). In a hypo-osmotic environment, fish eggs increase water intake to reach osmotic balance, and this water intake can increase the size of the eggs and lower their neutral buoyancy (Davenport et al. 1981; Thorsen *et al*. 1996). This allows them to remain in the water column at lower salinities, as seen in Baltic cod (*Gadus morhua*) populations (Nissling et al. 1994; Nissling and Westin 1997). Another adaptation to low salinity environment in the Baltic Sea is transitioning from pelagic to demersal spawning, as in the case of the flounder, where the demersally spawning Baltic flounder (*Platichthys solemdali*) has rapidly speciated from the sympatric pelagically spawning European flounder (*Platichthys flesus;* Momigliano et al. 2017; 2019, Nissling et al. 2017, 2002).

Since the 1980s, the Eastern Baltic cod (EBC) stock (SD24-SD32; (ICES 2021) has dramatically declined (Eero et al. 2023), and the body size at maturation has decreased from 40 to 20 cm (Casini et al. 2016; ICES 2021; Eero et al. 2023; Han et al. 2025). The declines in population and body size are mainly due to overfishing, but despite the complete fishing moratorium since 2019, the stock has been slow to recover (Eero et al. 2023; Han et al. 2025). Reduced oxygen, high infection load with liver parasites and low food availability in the Baltic Proper have been suggested to be connected to the slower growth as well (Eero et al. 2015, 2023; Casini et al. 2016; Ryberg et al. 2020; Orio et al. 2022). Increasing oxygen depletion has further reduced the area suitable for spawning in the Baltic Proper (Köster et al. 2003, 2005; Hinrichsen et al. 2016). Large volumes of water suitable for spawning currently occur mainly in the Bornholm Basin (SD25) in the southwest, but sometimes also in the Gdansk Basin and more infrequently in the Gotland Basin (SD 25, SD26 and SD28) (Köster et al. 2005; Casini et al. 2016; Bergström et al. 2025). Recently, suitable volumes of water for spawning have also been suggested to be found in the northern Baltic proper (SD27, 29 and 32; Bergström et al. 2025).

The cod in the Åland Sea (SD29), just north of the Baltic Proper, are larger and healthier than the cod in the southern Baltic (Figure 1) (Heimbrand et al. 2023; Bergström et al. 2025; Heimbrand and Limburg 2025). While most of these fish are in spawning condition during the breeding season (Bergström et al. 2025; Heimbrand and Limburg 2025), it is unclear whether they can successfully reproduce in the Åland Sea and nearby areas, or if the area is supported by drift of larvae and migration of young fish from known spawning areas further south in the Baltic Proper (Orio et al. 2019). In these northern areas, environmental salinity (5-10 psu) appears too low for pelagic development of EBC eggs and young larvae (Nissling and Westin 1991b). Interestingly, some Åland cod otoliths bear unique boron signatures, possibly suggesting a nursing area different from the cod in the southern Baltic Sea (Heimbrand and Limburg 2025). The uncertainty whether the Åland cod represents a self-sustaining population and is distinct from cod from the established spawning areas limits the development of effective management and conservation strategies for cod in the area.

**Figure 1:**
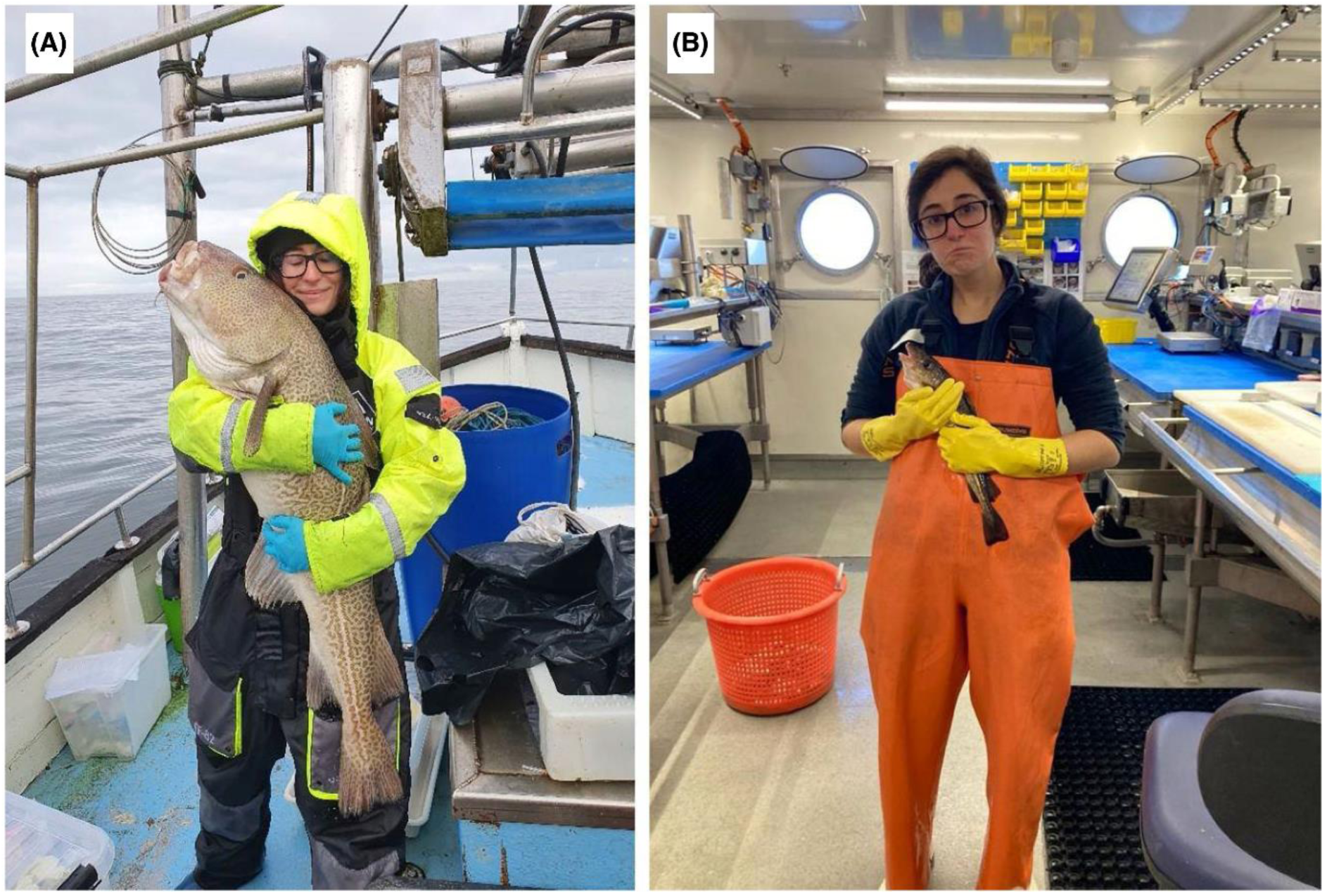
Comparison of northern Eastern Baltic cod (EBC) captured (A) in the Åland Sea (subdivision 29) versus (B) in the southern Baltic Sea (subdivision 25). MGO appears in both pictures for scale. Source: Heimbrand Y. and Lundberg K. E. (2025). Marin and Coastal Fisheries Vol 17 No. 4, 1-8. Photo credits: Y. Heimbrand (A) and O. Löfgren (B).

In this study, we investigated whether northern EBC (from the Åland Sea, SD29N) differs from southern EBC (from SD 25, 26 and 28) in its early life stages. If northern cod is locally adapted, we expect its eggs to show higher survival at low salinities found in the Åland Sea (5-7.5 psu), in Lågskär Deep (8-9 psu; southern Åland Sea) and southern SD29 (9-10 psu). We also investigated the possibility that Åland cod has evolved a spawning strategy suitable for a low-salinity environment. This could be achieved by larger and more buoyant eggs and larvae (Nissling et al. 1994; Nissling and Vallin 1996) or by eggs negatively buoyant and able to develop on the bottom, similar to the Baltic flounder (Momigliano et al. 2017). Specifically, we compared egg and larval survival at different salinities (7, 9.5 and 17 psu), egg and larval buoyancy, and survival on sediment in a series of laboratory experiments. This study is a continuation of Bergström et al. (2025), who studied fertilisation and egg development on northern cod, but included only fish *in situ* from the Åland Sea (5-7.5 psu), while here we kept northern and souther parental fish at the same salinity (17 psu) for at least 6 months and performed experiments on eggs and larvae laboratory conditions. In this study, we will refer to the northern and southern EBC as different populations due to their geographic and environmental divergence, even though the amount of reproductive isolation is not known.

## Material and Methods

### Broodstock acquisition, spawning and egg collection

Adult southern (eastern Baltic) cod (n = 130; 3:1 males and females; mean body length (cm) ± SE: 52,6 ± 1) were caught by trawl (Baltic International Trawl Survey - BITS) in ICES SD 25, 26 and 28 (Figure 2) in November 2021 (BITS-Q4), February 2022 BITS-Q1 and February 2023 (BITS-Q1). Adult northern (eastern Baltic) cod (n = 7; 3 males and 4 females; mean body length (cm) ± SE: 58,6 ± 3,6) were caught in the Åland Sea (SD 29) using gill nets near Grisslehamn on the eastern coast of Sweden in October-November 2023. This locationis ∼ 400 km north of the southern locations (∼ 250 km from SD28, ∼ 400 from SD 26 and ∼ 550 km from SD25 locations) and has much lower salinity decrease (5-7.5 psu; Figure 2) than the southern areas (13-17 psu; Figure 2). The fish were brought to the research station Ar in northern Gotland, Sweden (Figure 2), and distributed in three circular tanks (one for northern and two for southern cod) of 14 m^3^ (diameter 3.5 m, depth 1.46 m) water volume and supplied with air-saturated recirculated water (7 °C, 17 psu). Each tank was connected to a water treatment unit made up of a drum filter, UV filter, biofilter, trickle-filter, protein skimmer and heat exchanger for temperature regulation. Ozone was automatically dosed in the protein skimmer if the ORP values were below 290 mV. Natural brackish water (7 psu) was obtained from 35 m depth and filtered by a 0.5 μm particle filter and UV filter. In both tanks, salinity was adjusted to 17 psu by using artificial sea salt (Hoss, fish salt, Karlslunde, Denmark). The light regime was set to follow the natural light regime in Visby, Gotland (57°37′44″N 18°18′26″E) with Philips’s aquaculture light system 125W, including simulation of dawn and dusk. During the experimental period (May-June 2024), the fish were fed shrimp ad libitum every second day. Fish started to spawn in late May and were allowed to spawn freely. Fertilised eggs were collected daily (each day of collection was considered a batch) from the broodstock tanks, using one floating egg-collecting device (Cortney Ohs et al., 2019), per tank. The study was conducted under permits C 139/13 and 5.8.1810 169/2019 issued by the Ethical Committee on Animal Experiments in Uppsala, and 212-2021 and 03074-2022 issued by the Ethical Committee on Animal Experiments in Linköping.

**Figure 2:**
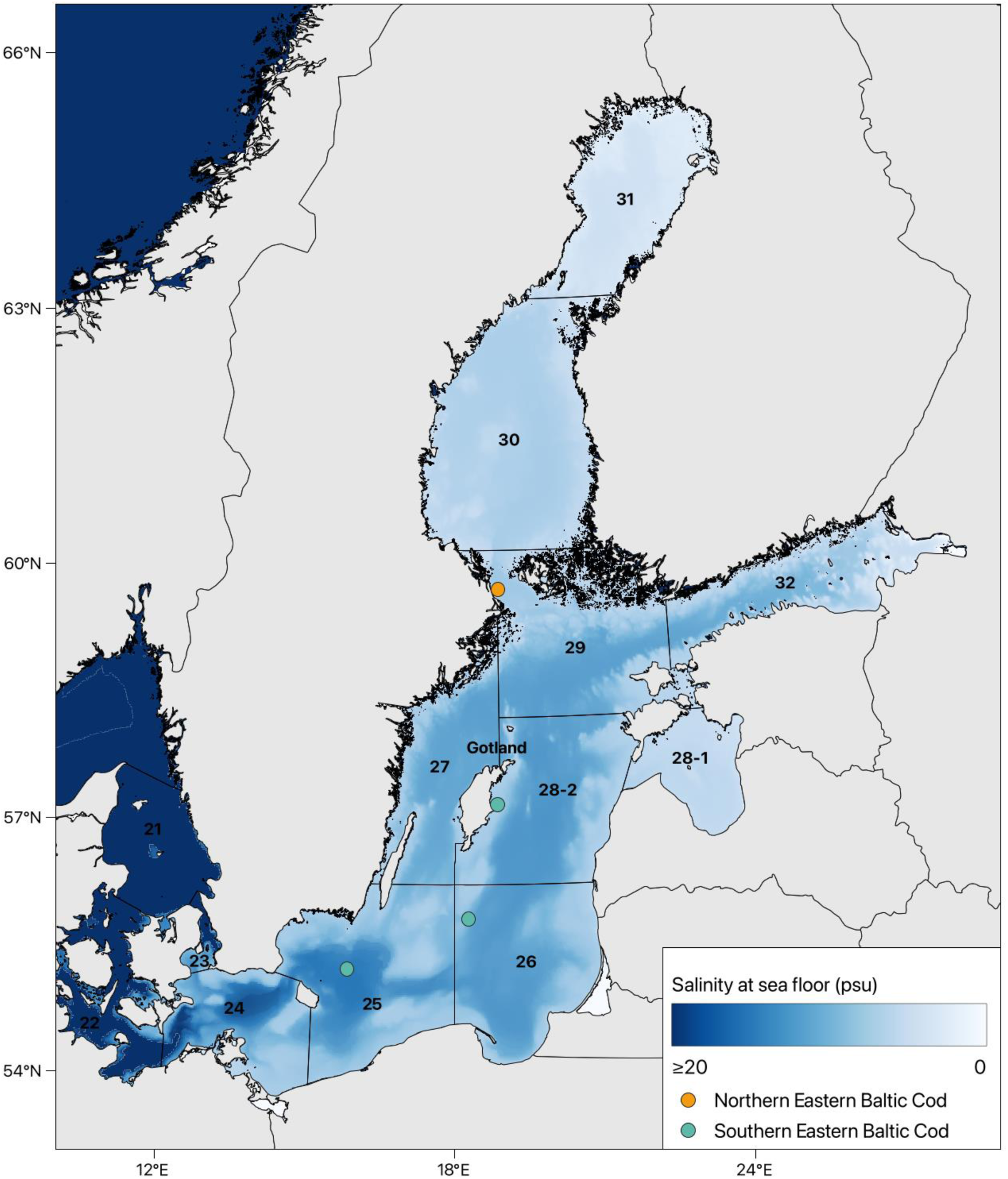
Geographical locations and salinity (psu) at the sea floor of the areas where the northern Eastern Baltic cod and southern Eastern Baltic cod were caught. Dots show the mean location where fish were caught in each section. Map generated using QGIS software using data on ICES areas (ICES Spatial Facility, ICES, Copenhagen) and data on salinity at the sea floor (E.U. Copernicus Marine Service Information of Baltic Sea Physics Reanalysis; https://doi.org/10.48670/moi-00013).

### Population differences in egg survival and hatching success at different incubation salinity treatments

We used eggs from six batches of northern cod and five batches of southern cod, collected between May 25 and June 19. We added 100 fertilised eggs to a 400 ml beaker filled with 7 °C water. We used three salinities: 7, 9.5 and 17 psu corresponding to the conditions at the southern and northern Baltic proper. Each salinity:population combination was triplicated within each batch. Egg developmental stage and quality were checked visually beforehand to ensure high quality of the batch and development of less than 20 h (zygote or cleave stages; (Hall et al. 2004). Two-thirds of the water was changed every second day, and dead eggs and larvae were discarded. Salinity and temperature were measured daily with an electronic conductivity meter (WTW Multi 3410 with TetraCon925 ids sensor, WTW Wissenschaftlich-Technische Werkstätten GmbH, Weilheim, Germany) and oxygen content was measured with 0.5 % accuracy using an oxygen probe (WTW FDO® 925 Optical IDS dissolved oxygen sensor) connected to the conductivity meter. Survival was assessed by counting the number of living larvae on day 2 after hatching, which for EBC larvae is 15 days after egg incubation.

### Egg size and neutral buoyancy

We estimated the neutral buoyancy of northern cod eggs using a salinity gradient (5-30 psu) column (Coombs 1981). Buoyancy was measured in a triplicate of 21-30 eggs from each of three daily batches collected between June 21 and 23. We used 5 glass density-balls with known and particular neutral buoyancy to estimate the salinity gradient in the column and the neutral buoyancy of each larva within each column (see (Garate-Olaizola et al. 2025) for details). Additionally, 36-44 eggs from each batch were photographed under a stereo microscope (NIKON SMZ800NI) and measured using IC Measure software. We used the average of three lines drawn across each egg as the individual egg diameter. We compared these measurements to the neutral egg buoyancy and size of cod from outside Gotland (SD 28; Nissling et al. 1994) and Bornholm Basin (SD 25; Garate-Olaizola et al. in prep.).

### Larval neutral buoyancy and size at different salinity treatments between populations

The larvae (2 days post-hatching) from the salinity experiment were used to measure neutral buoyancy and body length. Larval neutral buoyancy was assessed using the salinity gradient column method (Coombs 1981). Twelve to fifteen larvae were sedated using 0.05 mg/ml of benzocaine solution for five minutes, and allowed to sink for 10 minutes in the column before noting their position in the column. Fifteen larvae (or fewer if there were fewer survivors) from each replicate were photographed from a side using a stereo microscope (NIKON SMZ800NI), and their body length was later measured using IC Measure software.

### Egg survival and hatching success on sediment

Clay sediment was collected at the end of April 2024 on three locations east of Gotland (depths: 55, 58 and 60 m; Figure S1). The sediment was filtered to remove macrofauna and kept in a bucket with constant water inflow at the research Station Ar, for two months prior to the experiment. Before and during the experiment, the sediment was kept at 7 °C and 7 psu salinity and stirred 2-3 times per week.

The experiment was carried out using eggs from three batches of both northern and southern cod collected between the 4^th^ and 21^st^ of June. 100 mL of sediment was placed into a 400 mL beaker, followed by the addition of 300 mL of 13 psu water at 6 °C. We used an extra beaker as a control with 400 mL of 13 psu water at 6 °C but without sediment. One hundred eggs younger than 24 h were added to each beaker, and the absence of buoyant eggs in the beakers was confirmed visually. This setup was triplicated in each batch, and the beakers were incubated at 7 °C for 15 days, after which the number of hatched and living larvae was counted. Eggs that did not hatch died and remained on sediment. Water temperature, salinity and dissolved oxygen were measured on the first and last experimental days. No water change was done during the incubation to avoid sediment disturbance.

### Statistical analyses

We measured egg neutral buoyancy and egg size for northern cod for three batches. We examined variation in egg neutral buoyancy across batches by running a Generalised Linear Mixed Model (GLMM) from lme4 package (Bates et al. 2007) and the lmertest package extension (Kuznetsova et al. 2017), with the egg neutral buoyancy within each replicate as the dependent variable, batch as a fixed effect, and replicate as a random factor. We examined variation in egg size across batches by running a GLM (Generalized Linear Model) with egg size within as the dependent variable and batch as a fixed effect.

We analysed the neutral buoyancy of larvae with a Linear Mixed Model (LMM) run in lme4 package (Bates *et al*. 2015) and with the lmertest package extension (Kuznetsova et al. 2017) with population, salinity and their interaction as fixed effects, and batch within population and replicates within batch as random factors. Since the survival at 7 psu treatment was very low (very few survivors in northern cod), this treatment was excluded from the analysis. For body length, a GLMM was run with the mean larval body length in each replicate as a dependent variable and salinity, population and their interaction as fixed factors. Batch within population was included as a random factor in the model. Finally, we examined variation in survival between populations and between sediment treatments by analysing survival in each replicate using a glmmPQL with binomial distribution in the glmmTMB package (Brooks et al. 2017). We used the survival rate in each replicate as the dependent variable, and population, treatment, and their interaction as fixed effects. Batch within population and replicate within batch were added as random factors in the model.

In all the above models, statistical significance was computed using Anova function with Type III ANOVA with Satterthwaite’s method, performed using the car package (Fox and Weisberg 2019). When relevant, we tested pairwise differences between combinations of treatment factors with post-hoc comparisons performed using estimated marginal means (EMMs) with Tukey’s adjustment for multiple comparisons, implemented in the emmeans package (version 4.2.3; Lenth 2024). All analyses were performed using R v4.3.1 (R Core Team 2024).

## Results

### Larval survival at different egg incubation salinity treatments

Larval survival was significantly higher in the northern cod (χ^2^ = 748.34; p < 0.001; Table S1) at 17 psu and 9.5 psu salinity treatments (χ^2^ = 21.34; p < 0.001; Table S1; Table S2). The significant interaction term (χ^2^ = 147.05; p < 0.001; Table S1) indicated that survival was higher in northern and southern cod at 17 and 9.5 psu, but lower at 7 psu (Figure 3; Table S2). In both populations, larval survival was lowest at 7 psu treatment (Figure 3; Table S2).

**Figure 3:**
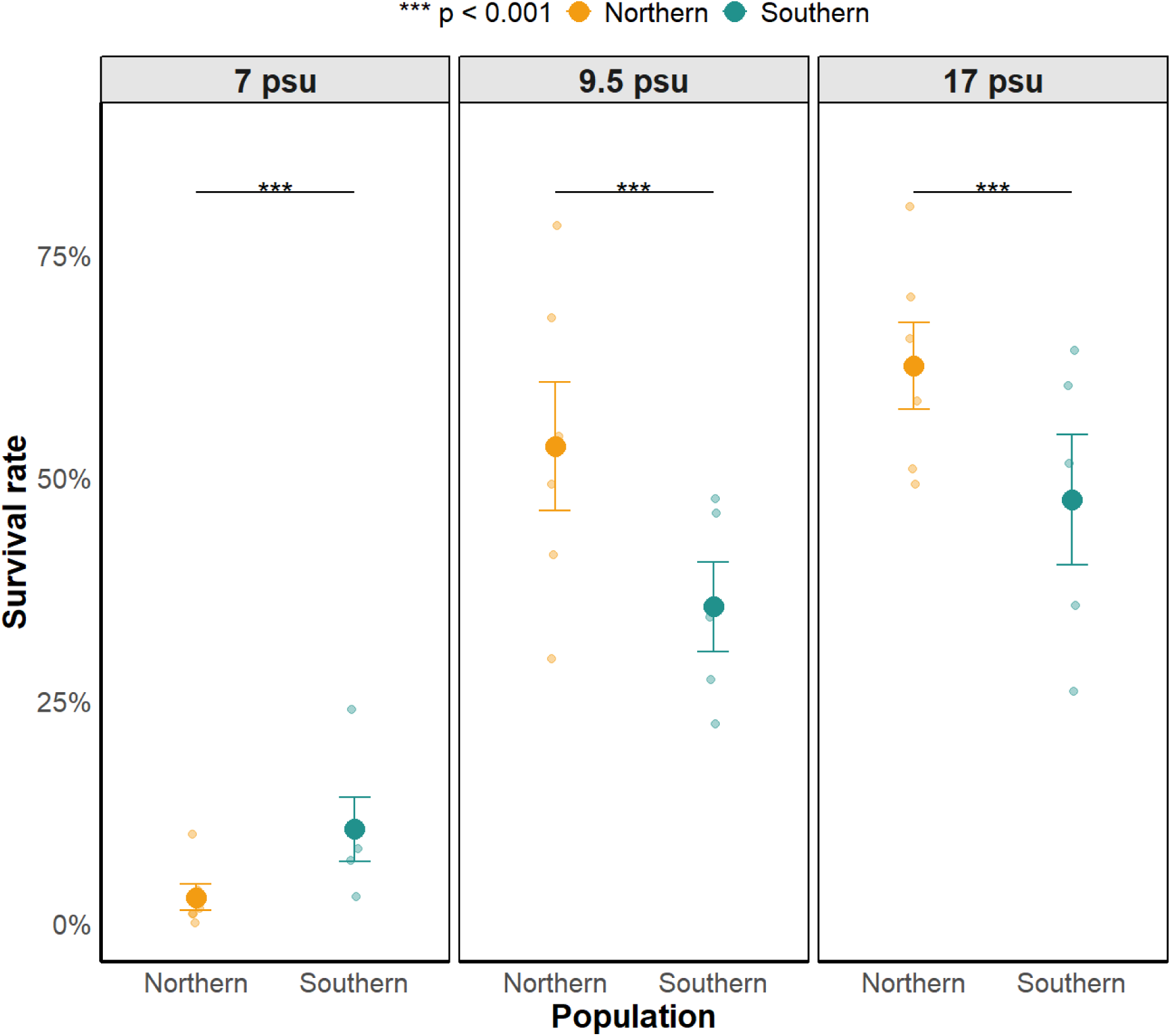
Survival rate of northern and southern eastern Baltic cod larvae incubated at different salinity conditions. Mean ± standard error (SE) showing larval survival in each salinity treatment (7 psu, 9.5 psu and 17 psu) for northern (orange) and southern (green) cod. Facets represent different salinity treatments. Survival is expressed as a percentage (%).

### Egg buoyancy and size in northern cod

Mean neutral buoyancy of northern cod eggs was 13.9 ± 0.06 psu, and ranged between 11.6 and 17.7 psu, including eggs from all three batches (Table 1). Eggs from batch 1 were significantly more buoyant than those from batch 2 and 3 (χ^2^ = 40.83; p < 0.001; Table S3). Average egg size was 1.67 ± 0.003 mm in diameter and ranged between 1.55 and 1.73 mm (Table 1), without significant variation between the batches (χ^2^ = 1.15; p = 0.563). For comparison, egg size and buoyancy for southern EBC are summarised in Table 1B (Nissling et al. 1994; Garate-Olaizola et al. in prep.).

**Table 1:** Summary of egg neutral buoyancy and size from this study for northern cod (SD 29) cod, for southern cod from SD 28, from (Nissling, Kryvi and Vallin 1994) and from SD 25, from Garate-Olaizola *et al*. manuscript in preparation.

| Cod population | Northern cod (SD29) | Southern cod (SD28) | Southern cod (SD25) |
| --- | --- | --- | --- |
|  | Egg neutral buoyancy |  |  |
| Number of eggs | 241 | NA | 890 |
| Mean (psu) ± SE | 13.9 ± 0.06 | 14.5 ± 1.2 | 14.9 ± 0.04 |
| Range (psu; min-max) | 11.6 – 17.7 | 12.3 – 18.3 | 12.3 – 19.2 |
|  | Egg size |  |  |
| Number of eggs | 120 | NA | 1275 |
| Mean (mm) ± SE | 1.67 ± 0.003 | 1.66 ± 0.07 | 1.54 ± 0.002 |
| Range (mm; min-max) | 1.55 – 1.73 | 1.49 – 1.78 | 1.34 – 1.82 |
| Source | This study | (Nissling, Kryvi and Vallin 1994) | Garate-Olaizola *et al.* manuscript in preparation |

### Larval neutral buoyancy and size at different salinity treatments

We found a significant effect of salinity treatment on larval neutral buoyancy (χ^2^ = 163.79; p < 0.001; Table S4; Figure 4a), with lower environmental salinities resulting in more buoyant larvae (Figure 4a; Table S5). However, we did not find a significant difference between the populations (χ^2^ 61 = 0.45; p = 0.503) nor a significant interaction between population and salinity (χ^2^ = 1.62; p = 0.203; Table S4; Figure 4a). There was no significant variation in body length between populations or among salinity treatments (χ²₁, ₈₀ = 0.41, p = 0.521; χ²₂, ₈₀ = 0.97, p = 0.615; Table S6; Figure 4b). Similarly, the interaction between population and salinity was not significant (χ²₂, ₈₀ = 0.65; p = 0.722; Table S6).

**Figure 4:**
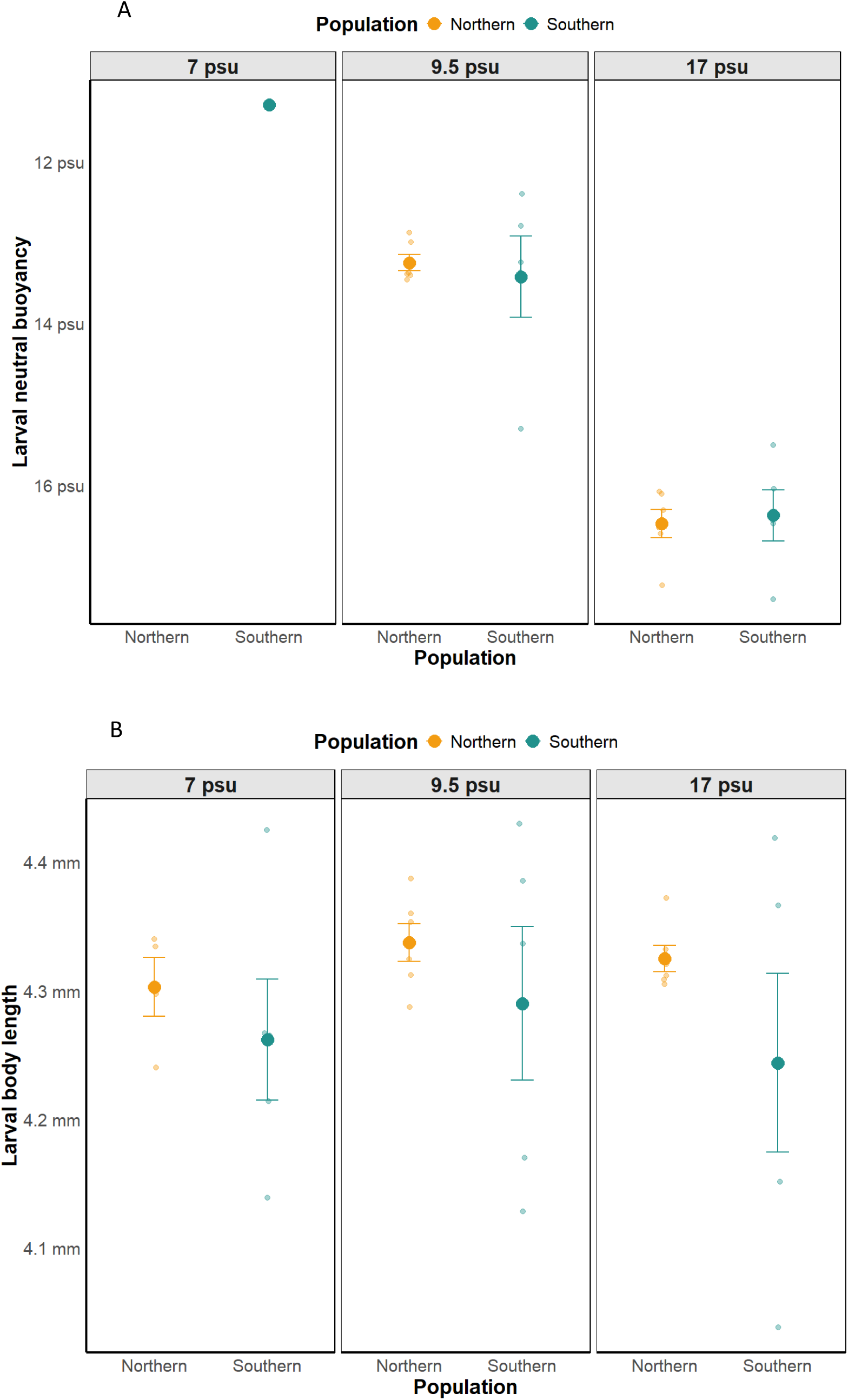
Larval neutral buoyancy (psu) and body length (mm) after incubation at different salinity conditions. Mean ± standard error (SE) of A) neutral buoyancy and B) body length of larvae incubated in 7, 11 and 17 psu).

### Egg and larval survival on sediment

Larvae from both populations hatched in the sediment treatment at 13 psu, with 57 % survival in the northern and 35% in the southern population (χ^2^ = 10.41; p < 0.001; Table S7; Figure 5). Survival in the sediment treatment was not significantly lower than in the non-sediment treatment (χ²₁, ₃₆ = 2.79; p = 0.095; Table S7; Figure 5). The interaction between population and treatment was not significant (χ²₁, ₃₆ = 0.07; p = 0.797; Table S7).

**Figure 5:**
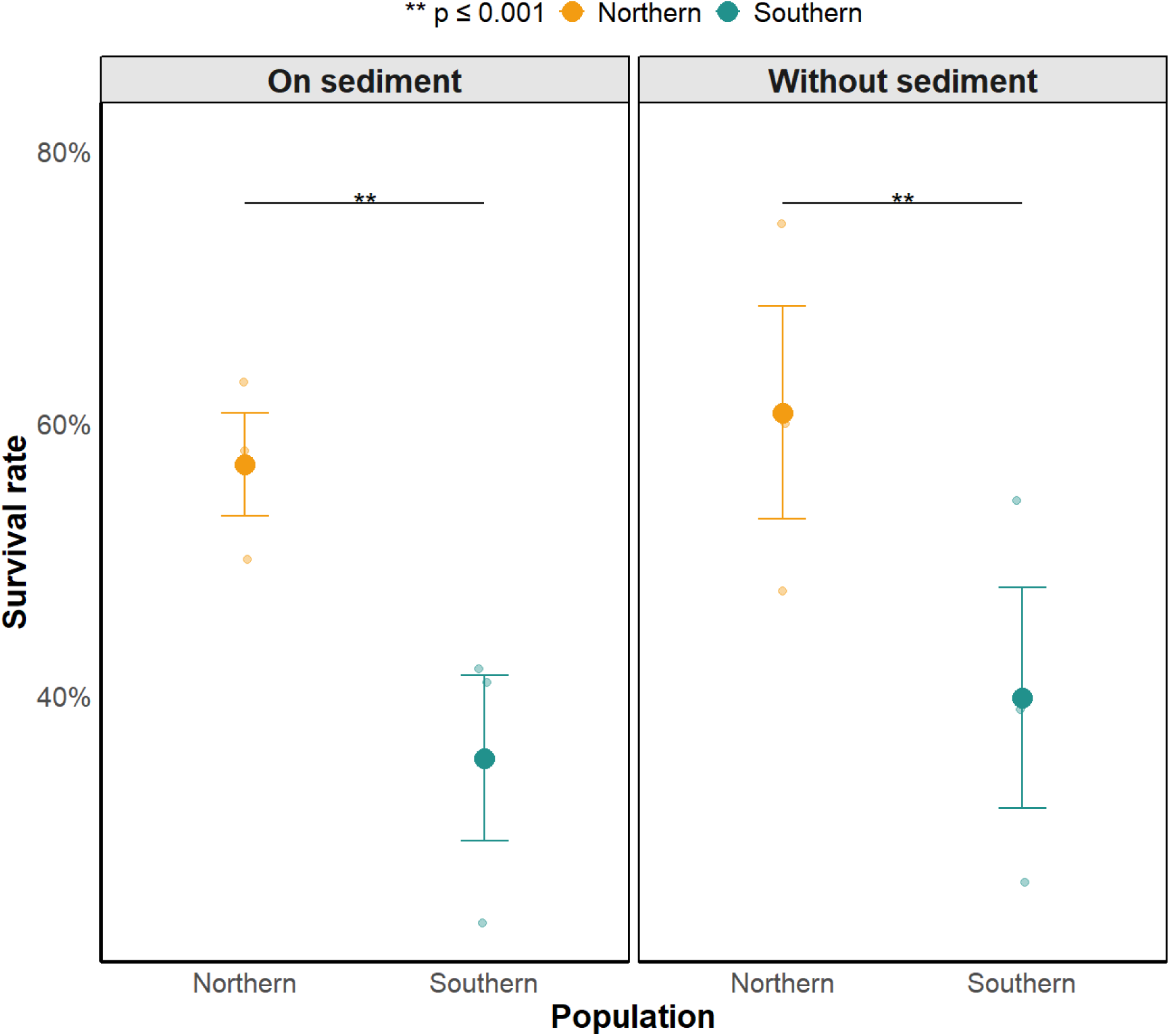
Survival rate of northern and southern cod eggs on sediment and non-sediment treatments. Mean ± standard error (SE) showing hatching success in each treatment (left sediment and right non-sediment) for northern (orange) and southern (green) cod. Survival is expressed as a percentage (%).

## Discussion

We investigated variation in survival and buoyancy between northern and southern cod at early life stages across a range of salinities and on sediment. Egg survival was notably low at 7 psu, indicating that successful egg development is unlikely in the Åland Sea. However, at intermediate salinity (9.5 psu), egg survival in northern cod was relatively high and higher than that in southern cod. Such conditions occur in southern SD29 (SD29S; northern Baltic Proper; Bergström et al. 2025) Furthermore, bottom salinity in the Lågskär Deep in the southern Åland Sea is also close to this value, ranging from 8 to 9 psu (Westerlund et al. 2022). However, both northern and southern cod required salinities >11 psu for egg and larval buoyancy, suggesting that they will sink to the bottom with the conditions prevailing in SD29. Our results also indicate that eggs may develop and hatch on sediment, with no significant difference from the non-sediment treatment. These findings, together with those presented by Bergström et al. (2025) showing that egg fertilisation can occur at 9.5 psu (although fertilisation success is low), suggest that there is potential for successful spawning outside the known spawning grounds in the southern Baltic Proper.

### Comparing between-population survival of eggs and larvae at different salinity treatments

We found that eggs survival in both northern and southern populations decline from 17 to 9.5 psu, with particularly low survival at 7 psu. These results are in line with Bergström et al. (2025) for northern cod, and by Nissling and Westin (1991b) for southern cod (SD 28). Adaptation to low salinity is associated with shifts in the internal osmolarity, optimal salinity range and tolerance of eggs and larvae, aligned with local environmental salinity (Holliday and Jones 1967; Holliday 1969; Nissling et al. 1994; Agarwal et al. 2024). In this context, higher tolerance to low salinity denotes the ability to maintain relatively higher early life survival at reduced salinity environment, as reported for EBC as compared to Western Baltic and Atlantic cod (Westin and Nissling 1991; Nissling et al. 1994; Nissling and Westin 1997). Here, we found that northern cod did not exhibit higher survival at the lowest salinity than the southern EBC population, indicating a lack of adaptation to the conditions in the Åland Sea (5-7.5 psu) at early life stages. However, our results indicate that over 50% of the northern larvae survived at 9.5 psu, which is significantly higher than what we found in the southern EBC. Successful hatching at 9.5 psu, combined with recent findings of well-oxygenated 9 psu conditions in SD29S and Lågskär Deep, may suggest that spawning is feasible outside the previously known grounds. Our study excluded fertilisation at 7.5 and 9 psu, which can be a major factor restricting successful spawning by limiting sperm motility. However, Bergström *et al*. (2025) found that fertilisation and early embryonic development are limited but can occur at 9 psu, supporting the idea that EBC might be able to successfully spawn in SD29S and Lågskär Deep.

### Egg buoyancy and egg size

Our results show that northern EBC eggs require ∼14 psu salinity to reach neutral buoyancy. These findings are consistent with previous results from the Åland Sea (Bergström *et al*. 2025). We also found that northern and southern cod eggs are similar with respect to buoyancy and size, our results being within the range of those described earlier for EBC (10.9-19.2 psu for neutral buoyancy and 1.37-1.82 in diameter; Nissling et al., 1994; Garate-Olaizola et al. in prep). Other pelagic spawners present in the Baltic Proper, such as plaice (*Pleuronectes platessa*) and European flounder, show similar values ranging from 14 to 17.7 psu and from 11.8 to 16.7 psu, respectively (Nissling et al. 2002; Nissling et al. 2017). These are lower values than those of their conspecifics in the western Baltic Sea (SD 24; Nissling *et al*. 2002), reflecting adaptation to a low-salinity environment. Similarly, northern and southern cod eggs were more buoyant and larger than eggs from western Baltic cod (Nissling and Westin 1997), and particularly more buoyant and larger than eggs from Atlantic cod (Solemdal and Sundby 1981), reflecting specific adaptation to the Eastern Baltic Sea environment. However, there was no difference in egg size and buoyancy between northern and southern EBC, indicating that the northern cod are not adapted to lower salinity conditions by increased egg buoyancy.

Demersal spawning, a common strategy in freshwater and low-salinity environments, is an alternative adaptation to low salinity (Orton 1957; Baras et al. 2018; Chen et al. 2021). The rapid speciation of Baltic flounder from the European flounder in the central and northern parts of the Baltic Sea by evolution of a demersal spawning strategy (Momigliano *et al*. 2017, 2019; Nissling *et al*. 2017) is an intriguing example of adaptive divergence. In contrast to flounders, northern cod eggs were neither smaller nor less buoyant than those from the southern area, requiring ∼ 14 psu to be buoyant. Given that northern cod eggs can tolerate the salinity in Lågskär Deep and southern SD29, these results suggest that successful reproduction would require eggs to develop on the sediment.

### Larval neutral buoyancy and size at different salinity treatments between populations

This study is the first to compare traits of larvae originating from northern and southern EBC. Notably, our results show no difference in neutral buoyancy between northern and southern cod larvae, and we also found no difference in larval body length between populations or across salinity treatments. Other studies have reported within-species differences in relation to the salinity at which larval cod reach neutral buoyancy (Westerberg 1994; Nissling and Vallin 1996; Saborido-Rey et al. 2003; Hüssy 2011), as well as in larval size after hatching. For example, EBC larvae are more buoyant and smaller than Western Baltic cod, and more buoyant than Atlantic cod (Solemdal and Sundby 1981; Nissling et al. 1994; Westerberg 1994; Saborido-Rey et al. 2003; Hüssy 2011). Larval neutral buoyancy is associated with egg buoyancy and influenced by maternal effects (Saborido-Rey et al. 2003; Govoni and Forward Jr 2008). In this context, our results do not support the hypothesis that northern cod larvae are better adapted to lower-salinity conditions than southern cod.

We found that larval neutral buoyancy decreases with decreasing environmental salinity. This pattern aligns with previous studies on EBC, which reported a comparable salinity-dependent decline in larval buoyancy (Nissling and Vallin 1996; Schmidt et al. 2024; Garate-Olaizola *et al*. 2025). At the early larval stage, osmoregulation is passive, and the internal osmolarity changes together with the ambient salinity, affecting larval neutral buoyancy (Holliday 1969; Sclafani et al. 1997; Saborido-Rey et al. 2003). Changes in internal osmolarity likely occur within the subdermal space of larvae, which expands at low salinity and enhances buoyancy independently of larval size (Nissling and Vallin 1996). While our data reflect the expected osmotic adjustment to salinity, larvae were still negatively buoyant at 7 and 9.5 psu, indicating that in low-salinity sites such as Lågskär Deep and southern SD29 larvae would remain demersal unless they swim actively.

### Egg survival on sediment

We found that EBC eggs survive well on sediment in the laboratory, challenging the common assumption that cod eggs die when in contact with sediment (Hinrichsen et al. 2016). Our results open up the possibility that Baltic cod may successfully spawn via demersal spawning and negatively buoyant eggs in the Baltic proper apart from the historical spawning grounds, provided that oxygen levels and salinity (> 9 psu) are appropriate and fertilisation can occur. We found that survival on sediment compared to the non-sediment treatment was similar in both populations, suggesting this is not a particular strategy developed by northern cod. Egg survival on sediment could occur in the southern SD29 between the Åland Sea and Gotland Deep (Bergström et al. 2025). Interestingly, also the Lågskär Deep in the southern Åland Sea could also occasionally offer conditions enabling some successful reproduction. However, our laboratory conditions differ from natural environments in several aspects, including predation pressure and microbial composition of the sediment, which may influence survival in situ in the natural environmental conditions (DeBlois and Leggett 1993; Hansen and Olafsen 1999). Another key question is the amount of prey available for the larvae hatching at the bottom, since cod larvae feed on zooplankton found in the water column (Van der Meeren and Naess 1993; Zuzarte et al. 1996). Further research using mesocosm experiments and field studies will help to better understand how EBC eggs and larvae perform at the bottom of the Baltic Proper. Nevertheless, dispersal processes remain important, and the larval drift from SD28 and SD26, alongside adult and juvenile migration, can be considerable (Bergström et al. 2025). Further research is needed on migratory patterns, nursery areas, and genomic divergence to better understand EBC population structure and explain the occurrence of healthy cod in the Åland Sea.

Due to logistic reasons, we had much fewer parental fish from northern cod than from southern cod. The parental fish from the two areas were kept in separate, single tanks. While temperature and salinity were similar in both tanks, we cannot rule out that our results are influenced by tank-specific effects affecting egg quality prior to spawning. Furthermore, as larval size is influenced by maternal effects (Marteinsdottir and Steinarsson 1998; Nissling et al. 1998; Marteinsdottir and Begg 2002; Saborido-Rey et al. 2003), the fact that we had fewer northern fish is likely to explain the lower variation we found in some traits (especially in larval body size) in northern cod.

## Conclusions

Northern and southern EBC had similar egg size and egg and larval buoyancy, thus not displaying adaptive divergence in these traits. However, the significantly higher survival of northern cod eggs in 9.5 psu compared to southern EBC, and high survival on sediment, are potential signs for successful reproduction in the northern Baltic Sea. These observations call for further research such as mesocosm experiments testing for demersal egg development in natural conditions, and field surveys for the occurrence of eggs and larvae in areas of SD29 where salinity (9-10 psu) and oxygen conditions would potentially allow demersal development. Addressing this issue is particularly important for the management of the EBC, given the currently deteriorated status of the stock (ICES 2021; Bryhn et al. 2022; Eero et al. 2023).

## Acknowledgements

We thank Dick Tillberg for his help in collecting the fish in the Åland Sea. Alex Stockhaus, Guillaume Vigouroux and Olov Tiblom from NIRAS and Clara Lunderg, Emma Bevin and Lise Toll from Ox2 for coordinating and executing the sediment collection, Konstantinos Vaziourakis for providing information on how to properly store and keep sediment, Gunilla Rosenqvist for providing help with the facilities and communication for the sediment collection, and Baltic Waters Foundation for financial and logistic support and the ReCod staff for taking care of the facilities, of the eggs fish, and collecting the eggs before and during the experiment.

## Author contribution

Maddi Garate-Olaizola (Resources [Lead], Data curation [lead], Formal analysis [lead], Funding acquisition [equal], Investigation [lead], Methodology [equal], Project administration [lead], Visualisation [lead], Writing – original draft [lead], Writing – review & editing [lead], Validation [equal]), Johanna Fröjd Resources (Resources [lead], Project administration [equal], Writing – review & editing [equal], Validation [equal]), Vanda Larsson Åberg (Data curation [equal], Investigation [equal], Methodology [equal], Visualisation [equal], Writing – review & editing [equal], Validation [equal]), Alma Hodzic-Vázquez (Data curation [equal], Investigation [equal], Methodology [equal], Writing – review & editing [equal], Validation [equal]), Yvette Heimbrand (Resources [equal], Writing – review & editing [equal], Validation [equal]), Anders Nissling (Conceptualisation [lead], Funding acquisition [Lead], Writing – review & editing [equal], Validation [equal]), Jane W. Behrens Supervision [equal], Writing – review & editing [equal], Validation [equal]), Maria Cortázar-Chinarro Supervision [equal], Writing – review & editing [Lead], Validation [equal]), Ulf Bergström (Conceptualisation [lead], Funding acquisition [Lead], Writing – review & editing [equal], Validation [equal]), Anssi Laurila (Conceptualisation [lead], Funding acquisition [Lead], Supervision [Lead], Writing – review & editing [Lead], Validation [equal]).

## Data availability

The data underlying this article will be shared on request to the corresponding author.

