## Supplementary tables and figures for "Assessing local adaptation and divergence at early life stages within Eastern Baltic cod"

**Supplementary material**

**Figures**

**Figure S1: Picture from Google maps showing the three locations (in blue) where sediment was collected.** WGS 84 coordinates, latitude and longitude: Location P19: 57.38580, 19.60004; location P16; 57.53058, 19.64841 and location P20: 57.51989, 19.64693.

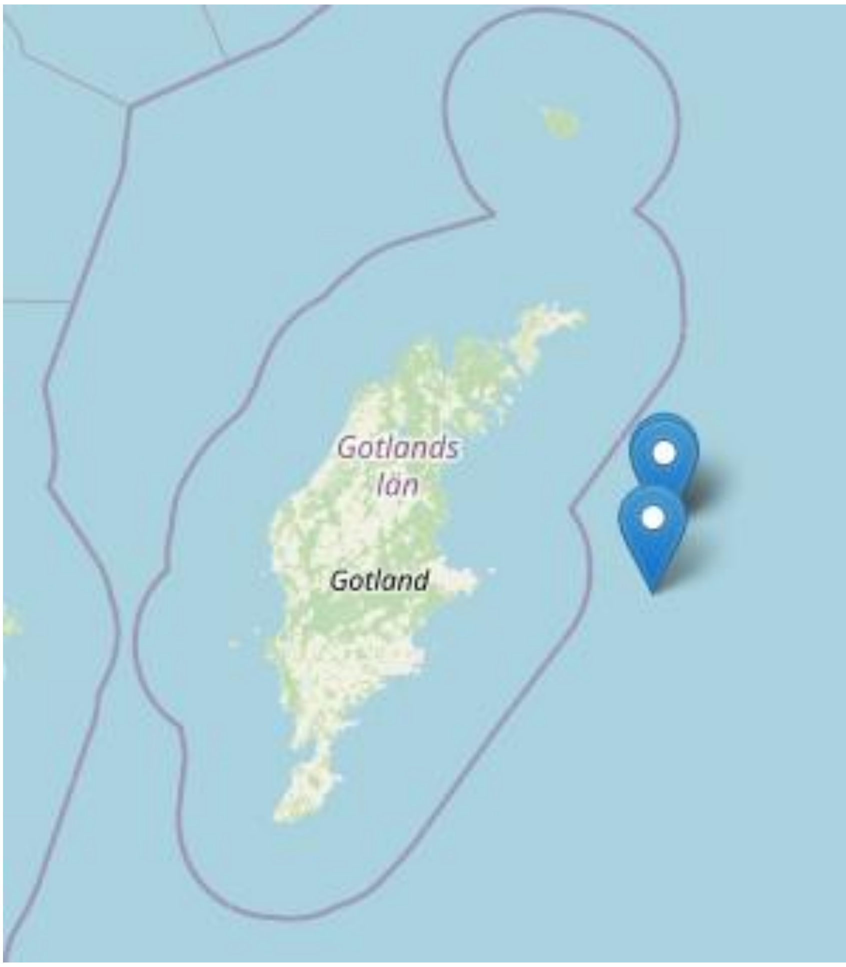

**Figure S2: Pictures of A) eggs developing on sediment in a beaker, with some already hatched larvae and B) a hatched larva from the sediment treatment.** A was taken with a Samsung Galaxy A42 and B with a stereo microscope (NIKON SMZ800NI) using IC Measure software.

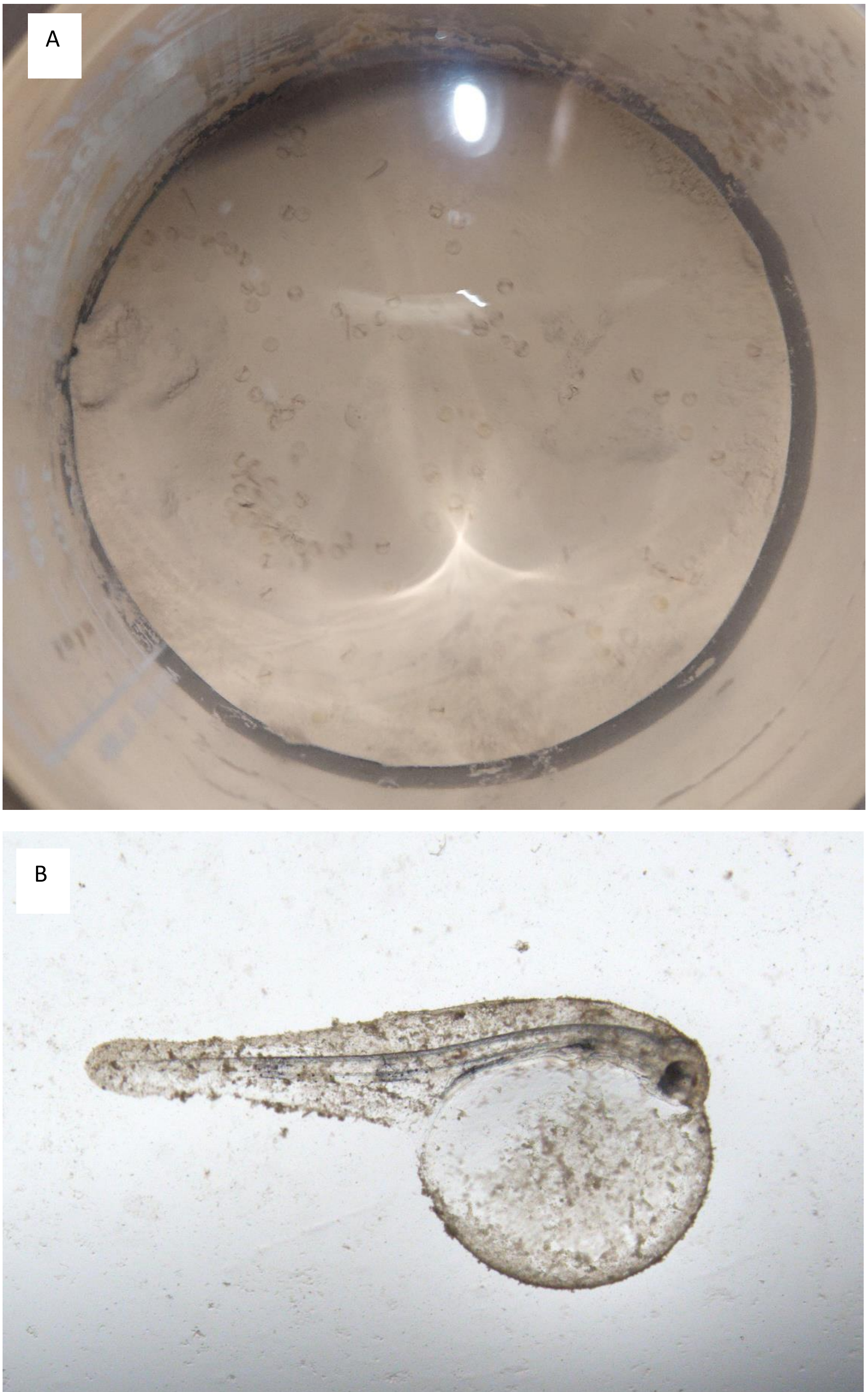

### Tables

**Table S1:** Results of the Generalized Linear Mixed-Effects Model Analysis testing the effect of salinity, population, their interaction, batch and its interaction with salinity on survival, including batches within populations and replicates within a batch as random factors. The table shows the $\chi^2$ test statistics ($\chi^2$), the denominator degrees of freedom (NumDF and denDF), and associated p-values (Pr (> $\chi^2$)) for the main effect and interactions, marked with * if significant.

| | $\chi^2$ | **NumDF** | **DenDF** | **Pr (> $\chi^2$)** |
|---|---|---|---|---|
| **Salinity** | 748.34 | 2 | 98 | < 0.001 * |
| **Population** | 21.32 | 1 | 98 | < 0.001 * |
| **Salinity:Population** | 147.05 | 2 | 98 | < 0.001 * |

**Table S2:** Results of pairwise post-hoc comparisons for the effects of population and salinity on survival using estimated marginal means (EMMs). The table presents the estimated differences (Estimate) between groups, standard errors (SE), test statistics (z.ratio), and adjusted p-values (p-value) based on Tukey's correction. Significant comparisons (p < 0.05) are marked with "*".

| | **Estimate** | **SE** | **z.ratio** | **P value** |
|---|---|---|---|---|
| **Northern - 17 psu Northern - 9 psu** | 0.38 | 0.07 | 5.35 | < 0.001 * |
| **Northern - 17 psu Northern - 7 psu** | 4.13 | 0.15 | 27.30 | < 0.001 * |
| **Northern - 17 psu Southern - 17 psu** | 0.65 | 0.14 | 4.62 | < 0.001 * |
| **Northern - 17 psu Southern - 9 psu** | 1.18 | 0.14 | 8.29 | < 0.001 * |
| **Northern - 17 psu Southern - 7 psu** | 2.77 | 0.16 | 17.70 | < 0.001 * |
| **Northern - 9 psu Northern - 7 psu** | -3.75 | 0.15 | -25.06 | < 0.001 * |
| **Northern - 9 psu Southern - 17 psu** | -0.28 | 0.14 | -1.96 | 0.367 |
| **Northern - 9 psu Southern - 9 psu** | 0.80 | 0.10 | -15.7 | < 0.001 * |
| **Northern - 9 psu Southern - 7 psu** | -2.40 | 0.16 | -15.41 | < 0.001 * |
| **Northern - 7 psu Southern - 17 psu** | 3.48 | 0.19 | 18.01 | < 0.001 * |
| **Northern - 7 psu Southern - 9 psu** | -2.95 | 0.19 | -15.24 | < 0.001 * |
| **Northern - 7 psu Southern - 7 psu** | -1.36 | 0.20 | -6.65 | < 0.001 * |
| **Southern - 17 psu Southern - 9 psu** | 0.53 | 0.08 | 6.87 | < 0.001 * |
| **Southern - 17 psu Southern - 7 psu** | 2.12 | 0.10 | 21.08 | < 0.001 * |
| **Southern - 9 psu Southern - 7 psu** | -1.59 | 0.10 | -15.70 | < 0.001 * |

**Table S3:** Results of pairwise post-hoc comparisons for the effects of variation in egg neutral buoyancy across batches using estimated marginal means (EMMs). The table presents the estimated differences (Estimate) between groups, standard errors (SE), test statistics (z.ratio), and adjusted p-values (p-value) based on Tukey's correction. Significant comparisons (p < 0.05) are marked with "*".

| | estimate | SE | DF | t.ratio | p value |
|---|---|---|---|---|---|
| Batch 1 Batch 2 | - 0.04 | 0.01 | Inf | -4.39 | < 0.001 * |
| Batch 1 Batch 3 | 0.02 | 0.01 | Inf | 1.81 | 0.17 |
| Batch 2 Batch 3 | 0.06 | 0.01 | Inf | 6.20 | < 0.001 * |

**Table S4:** Results of the Linear Mixed-Effects Model Analysis testing the effect of population, salinity and their interaction on larval neutral buoyancy (excluding 7 psu), including batches within populations and replicates within a batch as random factors. The table shows the $\chi^2$ test statistics ($\chi^2$), the denominator degrees of freedom (NumDF and denDF), and associated p-values (Pr (> $\chi^2$)) for the main effect and interactions, marked with * if significant.

| | **NumDF** | **DenDF** | $\chi^2$ | **Pr (> $\chi^2$)** |
|---|---|---|---|---|
| **Population** | 1 | 61 | 0.45 | 0.50 |
| **Salinity** | 2 | 61 | 163.79 | <0.001 * |
| **Population:salinity** | 1 | 61 | 1.62 | 0.20 |

**Table S5:** Results of pairwise post-hoc comparisons for the effects of population and salinity on neutral buoyancy using estimated marginal means (EMMs). The table presents the estimated differences (Estimate) between groups, standard errors (SE), test statistics (z.ratio), and adjusted p-values (p-value) based on Tukey's correction. Significant comparisons (p < 0.05) are marked with "*".

| **Contrasts** | **Estimate** | **SE** | **Df** | **t.ratio** | **p-value** |
|---|---|---|---|---|---|
| **Southern – 7 psu**<br>**Southern - 9.5 psu** | -2.33 | 0.68 | 53 | -3.44 | 0.010 |
| **Southern – 7 psu**<br>**Southern - 17 psu** | -5.34 | 0.67 | 53 | -7.99 | < 0.001 * |
| **Southern – 7 psu**<br>**Northern - 9.5 psu** | -2.15 | 0.67 | 53 | -3.23 | 0.018 * |
| **Bornholm – 7 psu**<br>**Northern - 17 psu** | -5.49 | 0.67 | 53 | -8.22 | < 0.001 * |
| **Bornholm – 9.5 psu**<br>**Bornholm - 17 psu** | -3.01 | 0.32 | 53 | -9.47 | < 0.001 * |
| **Southern – 9.5 psu**<br>**Northern – 9.5 psu** | 0.18 | 0.32 | 53 | 0.58 | 0.978 |
| **Southern – 9.5 psu**<br>**Northern – 17 psu** | -3.15 | 0.32 | 53 | 9.82 | < 0.001 * |
| **Southern – 17 psu**<br>**Northern – 9.5 psu** | 3.19 | 0.30 | 53 | 10.50 | < 0.001 * |
| **Southern – 17 psu**<br>**Northern – 17 psu** | -0.15 | 0.31 | 53 | -0.47 | 0.990 |
| **Northern – 9.5 psu**<br>**Northern – 17 psu** | -3.33 | 0.29 | 53 | -11.407 | < 0.001 * |

**Table S6:** Results of the Generalized Linear Mixed-Effects Model Analysis testing the effect of salinity, population and their interaction on body length, including batches within populations as random factors. The table shows the $\chi^2$ test statistics (Pr < $\chi^2$), the denominator degrees of freedom (NumDF and denDF), and associated p-values (Pr (> $\chi^2$)) for the main effect and interactions, marked with * if significant.

| | $\chi^2$ | **NumDF** | **DenDF** | **Pr (> $\chi^2$)** |
|---|---|---|---|---|
| **Salinity** | 0.97 | 2 | 80 | 0.615 |
| **Population** | 0.41 | 1 | 80 | 0.521 |
| **Salinity:population** | 0.65 | 2 | 80 | 0.722 |

**Table S7:** Results of the Generalized Linear Mixed-Effects Model Analysis testing the effect of population, treatment (sediment or no sediment) and their interaction on survival, including batches within populations and replicates within a batch as random factors. The table shows the $\chi^2$ test statistics ($\chi^2$), the denominator degrees of freedom (NumDF and denDF), and associated p-values (Pr (> $\chi^2$)) for the main effect and interactions, marked with * if significant.

| | $\chi^2$ | **NumDF** | **DenDF** | **Pr (> $\chi^2$)** |
|---|---|---|---|---|
| **Population** | 10.41 | 1 | 36 | 0.001* |
| **Treatment** | 2.79 | 1 | 36 | 0.095 |
| **Population:treatment** | 0.07 | 1 | 36 | 0.797 |
